# Mapping physiological ADP-ribosylation using Activated Ion Electron Transfer Dissociation (AI-ETD)

**DOI:** 10.1101/2020.01.27.921650

**Authors:** Sara C. Buch-Larsen, Ivo A. Hendriks, Jean M. Lodge, Martin Rykær, Benjamin Furtwängler, Evgenia Shishkova, Michael S. Westphall, Joshua J. Coon, Michael L. Nielsen

**Affiliations:** Proteomics program, Novo Nordisk Foundation Center for Protein Research, Faculty of Health and Medical Sciences, University of Copenhagen, Blegdamsvej 3B, 2200 Copenhagen, Denmark; Genome Center of Wisconsin, University of Wisconsin-Madison, Madison, WI, 53706, USA

## Abstract

ADP-ribosylation (ADPr) is a post-translational modification that plays pivotal roles in a wide range of cellular processes. Mass spectrometry (MS)-based analysis of ADPr under physiological conditions, without relying on genetic or chemical perturbation, has been hindered by technical limitations. Here, we describe the applicability of Activated Ion Electron Transfer Dissociation (AI-ETD) for MS-based proteomics analysis of physiological ADPr using our unbiased Af1521 enrichment strategy. To benchmark AI-ETD, we profiled 9,000 ADPr peptides mapping to >5,000 unique ADPr sites from a limited number of cells exposed to oxidative stress, corresponding to 120% and 28% more ADPr peptides compared to contemporary strategies using ETD and EThcD, respectively. Under physiological conditions AI-ETD identified 450 ADPr sites on low-abundant proteins, including *in vivo* cysteine auto-modifications on PARP8 and tyrosine auto-modifications on PARP14, hinting at specialist enzymatic functions for these enzymes. Collectively, our data provides new insights into the physiological regulation of ADP-ribosylation.

## INTRODUCTION

ADP-ribosylation (ADPr) is an emerging post-translational modification (PTM) regulating a variety of biological processes, including cell signaling and DNA damage repair, by modifying proteins with either one ADPr moiety (mono-ADP-ribosylation; MARylation) or several ADPr moieties (poly-ADP-ribosylation; PARylation). ADP-ribosyltransferases (ARTs) catalyze the modification by transferring ADPr units from NAD+ onto target proteins. The group of responsible enzymes can be divided into two groups depending on the conserved structural features; ARTCs (cholera toxin-like) and ARTDs (diphtheria toxin-like, better known as poly(ADP-ribosyl)polymerases, PARPs), of which the latter is the largest and most characterized group consisting of 17 members in human (Lüscher et al., 2017). Within the ARTDs, four and twelve enzymes are reported to possess PARylation and MARylation activity, respectively, while PARP13 is reported to be catalytic inactive (Lüscher et al., 2017).

The most thoroughly studied PARP enzyme is PARP1, which is predominantly investigated in relation to DNA damage repair (De Vos et al., 2012). Under genotoxic stress, PARP1 mainly modifies serine residues in conjunction with the co-factor Histone PARylation factor 1 (HPF1) (Bonfiglio et al., 2017b), with this induction abrogated by PARP inhibitors (Larsen et al., 2018). ADPr is a reversible PTM, allowing it to rapidly and situationally adapt to cellular stimuli in a timely manner. Two groups of erasing enzymes exist; the macrodomain fold and the dinitrogenase reductase-activating glycohydrolase (DraG) (Crawford et al., 2018). Poly(ADP-ribose) glycohydrolase (PARG) is one of the erasers containing the macrodomain fold, and is the key enzyme accountable for degrading PAR chains into MAR (D’Amours et al., 1999). Recently, ARH3 was reported to be a serine hydrolase belonging to the DraG group of enzymes (Crawford et al., 2018; Fontana et al., 2017).

In recent years, mass spectrometry (MS)-based proteomics methods have advanced as the primary approach for studying ADPr in a global and unbiased manner. As is the case with many PTMs, ADPr is very low abundant, making enrichment methods necessary. MS-based analysis is further complicated by the dynamic nature of ADPr, with a rapid enzymatic turnover of the modification (Alvarez-Gonzalez and Althaus, 1989), numerous modifiable amino acid residues (Altmeyer et al., 2009; Bonfiglio et al., 2017b; McDonald and Moss, 1994; Moss and Vaughan, 1977; Ogata et al., 1980; Sekine et al., 1989; Van Ness et al., 1980), and the highly labile nature of the modification (Bonfiglio et al., 2017a).

Several methods have been established for enrichment of ADPr-modified peptides for mass spectrometric analysis (Daniels et al., 2014; Gagne et al., 2018; Hendriks et al., 2019; Larsen et al., 2018; Zhang et al., 2013). For large-scale studies of ADPr, the two most widely used strategies are based either on affinity enrichment using either the Af1521 macrodomain, or on a combination of boronate-affinity enrichment coupled with chemical conversion of the ADPr modification into an amide using hydroxylamine. Whereas the Af1521-based enrichment strategy identifies the entire ADPr moiety on any amino acid residue (Larsen et al., 2018), the chemical-based strategy is only able to identify chemical marks on aspartic acid and glutamic acid residues (Li et al., 2019; Zhang et al., 2013). Moreover, while the Af1521 macrodomain is capable of identifying ADPr under endogenous conditions, knockdown of PARG is required to increase sensitivity for the chemical-based method, rendering investigation of ADPr under physiological conditions impossible (Li et al., 2019; Zhang et al., 2013).

We have previously demonstrated that MS acquisition relying on higher-energy collisional dissociation (HCD)-based fragmentation cannot be used for confidently localizing ADPr sites due to the labile nature of the bond between the modification and the amino acid residue (Larsen et al., 2018). Hence, the non-ergodic fragmentation propensity of electron transfer dissociation (ETD) fragmentation is necessary in order to confidently determine the exact amino acid residue being modified with ADPr. However, ETD is prone to non-dissociative electron transfer (ETnoD), resulting in fewer sequence-informative product ions (Ledvina et al., 2010), which we previously observed to especially affect ADPr-modified precursors (Larsen et al., 2018). To overcome the high degree of ETnoD, supplemental activation can be applied with electron-transfer higher-energy collisional dissociation (EThcD), which has been proven to be more effective than ETD for identification of ADPr sites (Bilan et al., 2017; Hendriks et al., 2019). However, EThcD did not have a major effect on the dissociation of charge-reduced ADPr peptide product ions (Hendriks et al., 2019).

By contrast, activated ion-ETD (AI-ETD) uses infrared photoactivation during the ETD reaction to overcome ETnoD, and has been shown to benefit glycoproteomics (Riley et al., 2019), top-down proteomics (McCool et al., 2019; Riley et al., 2018a; Riley et al., 2018b), as well as phosphoproteomics (Riley et al., 2017a). We therefore set out to investigate the utility of AI-ETD on an Orbitrap Fusion Lumos mass spectrometer for identification and confident localization of ADPr sites. Here, we explored different laser power settings for AI-ETD, and compared it to ETD and EThcD. Overall, AI-ETD proved very advantageous for analyzing ADPr-enriched samples, with AI-ETD mapping 79% and 23% more ADPr sites compared to ETD and EThcD, respectively. With the enhanced performance of AI-ETD, we could obtain a comparable depth of sequencing from markedly less input material. This increased sensitivity auspiciously enabled identification of 450 physiological ADPr sites, of which 60% were not previously identified under physiological conditions. Taken together, we demonstrate that AI-ETD is a superior dissociation technique for sensitive MS-based proteomics analysis of ADP-ribosylation, and facilitates analysis of physiological ADPr from limited amounts of starting material.

## RESULTS

### Evaluation of AI-ETD

In order to investigate the ability of AI-ETD to increase the dissociation of ADP-ribosylated peptides, we utilized our established Af1521 method that allows for efficient and high-purity enrichment of peptides modified by ADP-ribosylation (Figure 1A). As the binding affinity of the macrodomain is towards the terminal moiety of ADP-ribose (Karras et al., 2005), the Af1521 strategy is capable of detecting modification of any amino acid residue type by ADP-ribosylation (Larsen et al., 2018). For the purpose of initial optimization and direct comparison of AI-ETD with other dissociation methodologies, we purified ADP-ribosylated peptides from H_2_O_2_-treated HeLa cells. This sample was used as a technical standard for quadruplicate comparison of different dissociation types, with each experiment corresponding to a comparatively low amount of ADP-ribosylated peptides purified from only two million cells. We compared standard ETD to EThcD, which we previously demonstrated to be superior for the identification of ADP-ribosylated peptides (Hendriks et al., 2019). Additionally, we used AI-ETD at five different laser power settings, spanning the range of routinely used power settings (Ledvina et al., 2010; Riley et al., 2017b).

**Figure 1.**
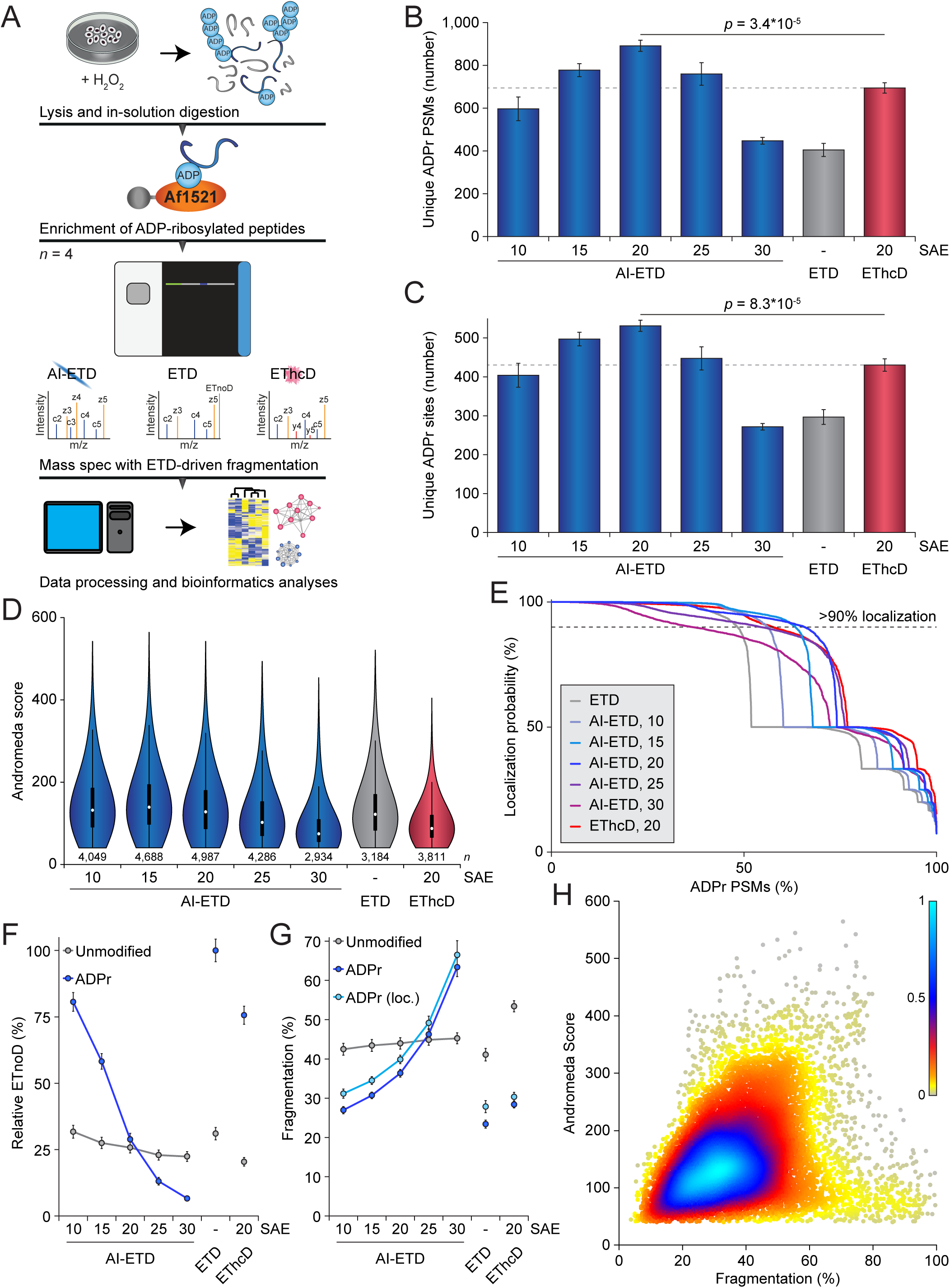
Comparison of AI-ETD vs. ETD and EThcD for mapping ADP-ribosylation. (A) Overview of the strategy employed to enrich ADP-ribosylated peptides. For technical comparison of dissociation methods, all samples were analyzed in technical quadruplicate; *n*=4. All dissociation methods were based on ETD to facilitate localization of the labile ADPr. (B) Overview of the number of ADPr peptide-spectrum-matches (PSMs) identified and localized (>90% probability) for each dissociation method and Supplemental Activation Energy (SAE). Error bars represent SD, *n*=4 technical replicates. (C) As **B**, but displaying the number of ADPr sites identified. (D) As **B**, but displaying the spectral quality (in Andromeda Score) of all identified ADPr-modified peptides. Distribution of data points is visualized, line limits; 1.5× interquartile range (IQR), box limits; 3^rd^ and 1^st^ quartiles, white dot; mean. Number of data points (*n*) is visualized below the distributions. (E) ADP-ribose localization probability plotted against the ranked fraction of all peptide-spectrum-matches (PSMs) resulting from the different dissociation methods and SAEs. Note that although all probabilities are displayed, only those over 0.9 were used for assignment of ADP-ribosylated peptides and sites. (F) Visualization of the average relative degree of non-dissociative electron transfer (ETnoD). Derived from all peptide-identified MS/MS spectra, and separately visualized for unmodified and ADP-ribosylated peptides. Error bars represent 5× SEM. (G) Visualization of the average degree of precursor fragmentation, calculated by dividing observed fragment ion peak intensity by the sum of non-ETD, ETnoD, and all fragment ion peak intensities. Derived from all peptide-identified MS/MS spectra, and separately visualized for unmodified, ADP-ribosylated, and localized ADP-ribosylated peptides. Error bars represent 5× SEM. (H) Spectral quality (in Andromeda Score) plotted against the average degree of precursor fragmentation, for localized ADP-ribosylated peptides. Coloring represents the relative density of dots in the plot, with higher values corresponding to higher density.

We observed a higher number of identified and localized unique modified ADP-ribosylated peptides for all five AI-ETD laser power settings, compared to pure ETD (Figure 1B and Table S1). The optimum energy setting was 20%, with a gradual fall-off in identification rate for both lower and higher laser power settings. As we described previously, EThcD outperformed pure ETD, but overall AI-ETD performed significantly better. AI-ETD identified 120% more ADPr-modified peptides than ETD, and 28% more ADPr-modified peptides than EThcD. When considering the number of unique ADP-ribosylation sites on proteins, we found the same trend (Figure 1C and Table S2), with AI-ETD mapping 79% and 23% more sites compared to ETD and EThcD, respectively.

In terms of spectral quality, as assessed by Andromeda Score, AI-ETD demonstrated the highest scores when using relatively low laser power settings, with an optimum at 15% (Figure 1D). Higher laser power settings of 25% and above, as well as EThcD, resulted in a decline in spectral quality, suggesting excessive peptide fragmentation under these circumstances. When considering localization of the ADP-ribose to specific amino acid residues within the peptide, we found that on average AI-ETD at laser powers 15% and 20%, as well as EThcD, significantly outperformed pure ETD (Figure 1E). However, when filtering ADPr for a stringent localization cut-off of 90%, AI-ETD at 20% laser power was superior.

Previously, we demonstrated complementarity of trypsin and Lys-C digestions for identification of unique ADPr sites (Hendriks et al., 2019). Thus, we purified ADPr peptides as described above, but using Lys-C instead of trypsin digestion. We subjected these peptides to technical quadruplicate analysis with a similar experimental setup, using the three optimal AI-ETD laser power settings. Overall, the aforementioned findings could be recapitulated, with the best performance of AI-ETD at laser power 20%, identifying 134% and 30% more ADPr peptides than ETD and EThcD, respectively (Figure S1A and Table S1). At the site-specific level, AI-ETD mapped 118% and 28% more ADPr sites than ETD and EThcD, respectively (Figure S1B and Table S2), and spectral quality was likewise improved by AI-ETD (Figure S1C).

Based on these observations, we wondered whether the increased performance of AI-ETD could be explained through increased dissociation of reduced charge-state precursors, i.e. decreased levels of ETnoD, and an overall more complete fragmentation of the ADP-ribosylated peptides. Intriguingly, AI-ETD considerably reduced the amount of ETnoD even at lower laser power settings, and dramatically reduced ETnoD to practically zero at the highest laser power settings (Figure 1F and S1D), whereas EThcD failed to resolve ETnoD product ions to the same extent. Concomitantly, the unmodified peptides within the same samples were only modestly affected by AI-ETD, as these were generally short peptides that did not display high levels of ETnoD to start with. Moreover, this highlights a striking susceptibility of ADPr peptides to AI-ETD. Correspondingly, when considering the overall level of precursor fragmentation, we found a notably increased fragmentation of ADPr peptides with increased levels of AI-ETD laser power, whereas unmodified peptides were not affected as much (Figure 1G and S1E). Contrarily, EThcD did not greatly increase fragmentation of ADPr peptides, but did affect unmodified peptides. Overall, AI-ETD reduced ETnoD and led to more complete fragmentation of specifically ADPr peptides.

ADPr peptides achieved a high level of fragmentation at the highest AI-ETD laser power settings, yet this did not result in the highest number of identifications. To investigate this, we compared the degree of peptide fragmentation to spectral score. For unmodified peptides, there was a global correlation with higher degrees of fragmentation corresponding to higher spectral quality (Figure S1F). For localized ADPr peptides, we found the highest quality spectra to frequently reside in the range from 30% to 45% fragmentation, and noted a strong decline in spectral quality when fragmentation exceeded 50% (Figure 1H). Indeed, when considering identified but non-localized ADPr peptides, a considerably higher number of these were >50% fragmented (Figure S1G), suggesting that very high degrees (>50%) of ADPr peptide fragmentation are detrimental to faithful identification and localization of this PTM. Collectively, we show that AI-ETD is superior to ETD and EThcD for confident profiling of ADPr sites.

### Benchmarking AI-ETD for in-depth profiling of ADPr

To gauge the performance of AI-ETD in the context of deep analysis of the ADP-ribosylome, we employed our previously described high-pH fractionation on ADPr-modified peptides purified from H_2_O_2_-treated HeLa cells (Figure S2A) (Hendriks et al., 2019; Larsen et al., 2018). Since we established that AI-ETD is more effective than ETD and EThcD, we used AI-ETD at the optimal laser power setting of 20% for all further experiments. Additionally, to address the large amount of starting material usually required for proteomic profiling of ADPr (Abplanalp et al., 2017; Hendriks et al., 2019; Larsen et al., 2018; Li et al., 2019; Zhang et al., 2013), we performed mass spectrometric analysis on a considerably smaller amount of sample. In total, 8,939 unique ADPr-modified peptides were identified (Table S1), mapping to 5,206 confidently localized ADPr sites (Figure 2A and Table S2). An intermediate total number of ADPr sites was identified compared to previous studies, falling just short of the current single-study record of 7,000 (Hendriks et al., 2019). Nevertheless, the relative amount of input material analyzed using AI-ETD was only ∼20%, yet still facilitated identification of ∼75% of the total number of sites (Figure 2A). Additionally, our AI-ETD profiling of the ADP-ribosylome from limited input revealed ADPr modification of 1,800 proteins (Table S3), corresponding to 84% of the number of proteins identified in the largest single MS study to date (Hendriks et al., 2019). More than half of the ADPr target proteins were modified on two or more distinct ADPr sites, demonstrating deep coverage of this low stoichiometry PTM (Figure S2A). Collectively, our benchmarking data exemplify the great sensitivity of AI-ETD.

**Figure 2.**
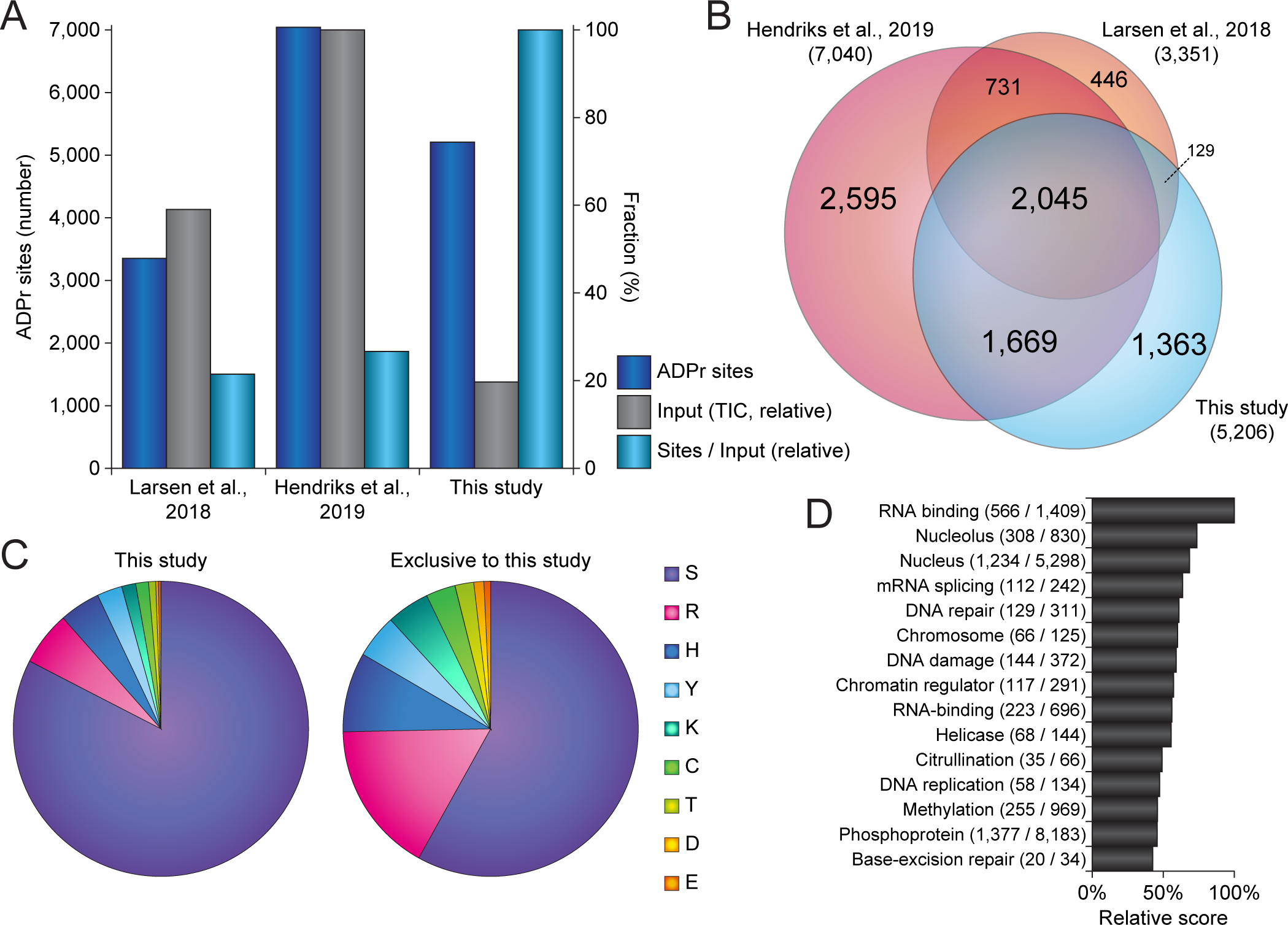
Benchmarking of AI-ETD. (A) Overview of the number of ADPr sites identified in this experiment, as compared to previous screens performed with the same ADPr-modified peptide enrichment strategy (Hendriks et al., 2019; Larsen et al., 2018). “Input” corresponds to the relative amount of the summed total ion current (observed at the MS1-level) across the analytical gradients of all experiments, and correlates with effective sample load on the column. (B) Scaled Venn diagram visualizing distribution of identified ADP-ribosylation sites in this experiment compared to two other ADP-ribosylation studies (Hendriks et al., 2019; Larsen et al., 2018). (C) Pie-chart overview of the amino acid residue distribution of all ADP-ribosylation sites identified in this experiment (left pie-chart), and ADP-ribosylation sites exclusively identified in this study (right pie-chart) as compared to two other ADP-ribosylation studies (Hendriks et al., 2019; Larsen et al., 2018). (D) Term enrichment analysis using Gene Ontology annotations and Uniprot keywords, comparing proteins identified to be ADP-ribosylated in this study to the total proteome. Relative score is based on multiplication of logarithms derived from the enrichment ratio and the *q*-value. All displayed terms were significant with *q*<0.02, as determined through Fisher Exact Testing with Benjamini-Hochberg correction. The full term enrichment analysis is available in Table S5.

Overall, 1,798 (35%) and 2,045 (39%) sites were previously described by either or both our other studies, respectively (Figure 2B). Moreover, we identified 1,363 (26%) previously undescribed ADPr sites (Table S4). In terms of amino acid residue specificity, we observed the large majority of modification to occur on serine residues (83%), followed by modification of arginine (5.9%), histidine (4.5%), and tyrosine (2.8%) residues (Figure 2C, left). When only considering the subset of ADPr sites exclusively identified in this study, we still found the majority of these to reside on serine residues (58%), but with a relatively larger fraction of ADPr also targeting arginine (17%), histidine (8.7%), tyrosine (4.8%), lysine (4.7%), cysteine (3.2%), threonine (2.1%), aspartic acid (1.1%), and glutamic acid (0.7%) residues (Figure 2C, right).

The H_2_O_2_-induced ADP-ribosylome profiled using AI-ETD largely resided on proteins with functional annotations that are well-known in the context of ADP-ribosylation biology, including predominant modification of nuclear, nucleolar, and chromatin-centric proteins, and modification of proteins involved in RNA binding, DNA repair, chromatin regulation, and DNA replication (Figure 2D and Table S5). Taken together, AI-ETD is congruous with deep analysis of the ADP-ribosylome from a limited pool of starting material.

### Improved analysis of physiological ADPr using AI-ETD

ADP-ribosylation is frequently investigated in a non-physiological manner, e.g. by genetic perturbation of ADP eraser enzymes (Daniels et al., 2014; Zhang et al., 2013), or by exposure of cells to high levels of genotoxic or proteotoxic stress, which both serve to boost cellular levels of ADPr in order to facilitate detection. Levels of ADPr are much lower under physiological conditions, e.g. in unperturbed cells growing under standard conditions. This makes physiological ADPr technically challenging to study, despite the biomedical importance of elucidating how ADPr is regulated at baseline conditions, e.g. in clinical samples or patient biopsies. Therefore, we set out to evaluate the ability of AI-ETD to profile the physiological ADP-ribosylome.

Using our established strategy, we purified ADPr-modified peptides from unperturbed HeLa cells growing under standard conditions. To minimize the requirement for mass spectrometric machine time we did not perform high-pH fractionation, and analyzed the physiological ADPr samples as single shots using AI-ETD. In total, we identified 703 unique ADPr-modified peptides, corresponding to 450 ADPr sites, residing on 295 proteins (Figure 3A). Using AI-ETD, we identified 79% and 26% more ADPr sites compared to single shot and high-pH fractionated measurements from our previous study (Larsen et al., 2018), respectively. When considering the best replicate, we were able to map 387 sites from a single run spanning 50 minutes of analytical gradient time, while on average AI-ETD allowed us to identify and confidently localize up to 8 physiological ADPr sites per minute of machine time.

**Figure 3.**
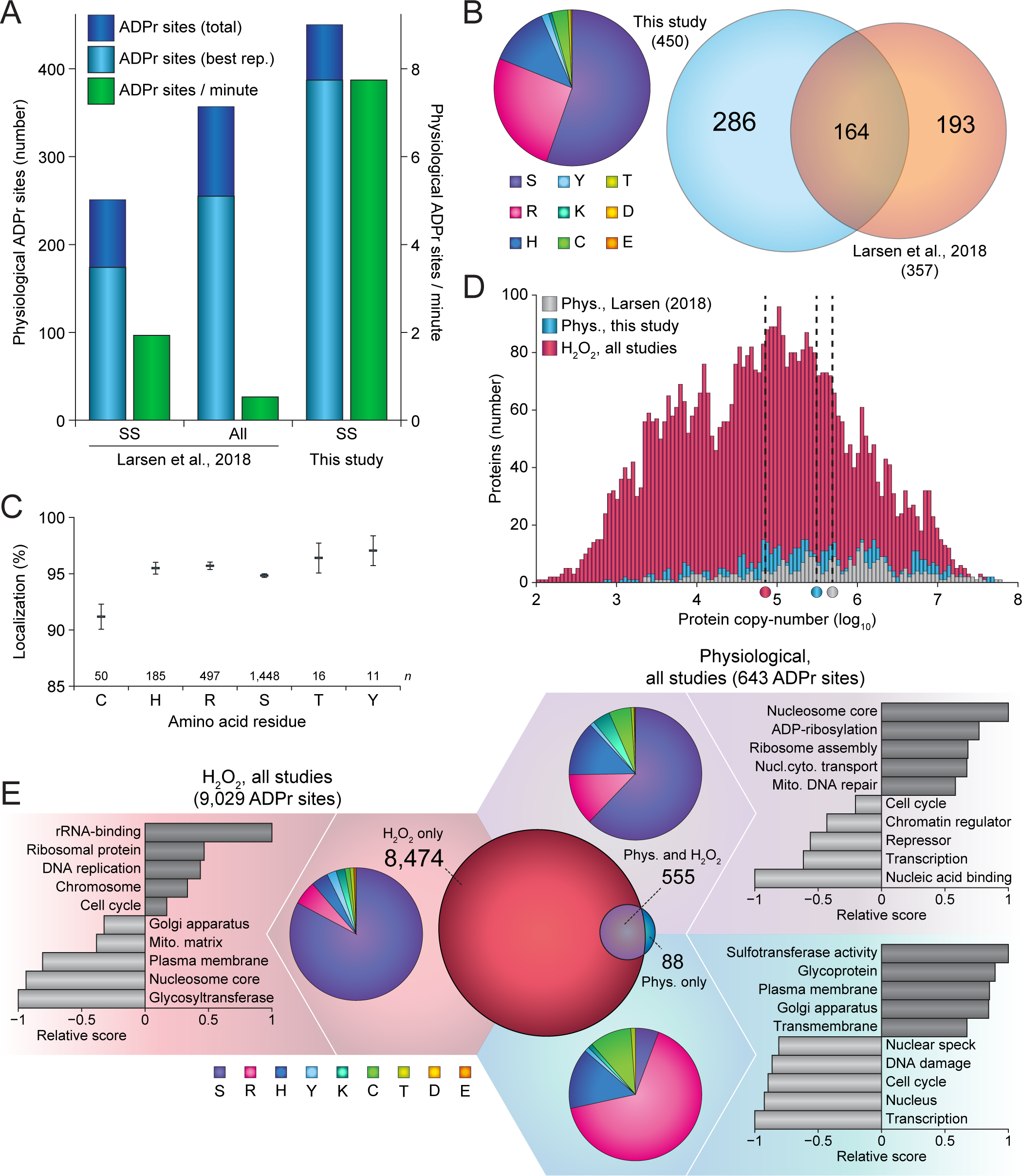
Analysis of physiological ADPr using AI-ETD. (A) Overview of the number of physiological ADPr sites identified in this study, compared to a previous study which also included physiological ADPr (Larsen et al., 2018). “Total” represents the number of unique ADPr sites identified across all four cell culture replicates, “Best rep.” represents the number of unique ADPr sites identified in the best replicate. “ADPr sites / minute” was calculated by dividing the best replicate number of ADPr sites by the effective analytical gradient time in minutes. “SS”; single-shot analysis, “All”; high-pH fractionated analysis. (B) Left; pie-chart overview of the amino acid residue distribution of all physiological ADP-ribosylation sites identified in this study. Right; scaled Venn diagram visualizing distribution of identified physiological ADP-ribosylation sites in this study compared to Larsen et al., 2018 (Larsen et al., 2018). (C) Average localization probability across all ADPr PSMs that were at least partially localized (>51%) to each specific amino acid residue type. Only amino acids with >10 partially localized PSMs were included. Error bars represent SEM, number of data points (*n*) is indicated. (D) Depth of sequencing analysis, plotting number of identified physiological ADPr target proteins versus their known copy-numbers. Proteins ADP-ribosylated in response to H_2_O_2_ were derived from this study, as well as various other studies (Bilan et al., 2017; Jungmichel et al., 2013; Larsen et al., 2018; Martello et al., 2016; Zhang et al., 2013). Protein copy-numbers were derived from a deep HeLa proteome study (Bekker-Jensen et al., 2017), which covers the vast majority (>99%) of all ADP-ribosylated proteins identified. Dotted lines with color indicator below represent the median protein copy-number for ADPr target proteins within each respective subset. (E) ‘Fidget spinner’ analysis, comprised of; scaled Venn diagram (center) visualizing distribution of H_2_O_2_-induced and physiological ADP-ribosylation sites; pie-chart overviews (around the center) of the amino acid residue distribution of each subset of ADPr sites, as indicated by background coloring; term enrichment analyses (outermost graphs) using Gene Ontology annotations and Uniprot keywords, comparing proteins identified to be ADP-ribosylated in each subset of ADPr sites as compared to the other subsets, indicated by background coloring. Relative score is based on multiplication of logarithms derived from the enrichment ratio and the *q*-value. All displayed terms were significant with *q*<0.02, as determined through Fisher Exact Testing with Benjamini-Hochberg correction. The full term enrichment analysis is available in Table S5. Background and Venn coloring, red; H_2_O_2_-only, purple; both H_2_O_2_ and physiological, blue; physiological only.

Out of the 450 ADPr physiological sites we identified, 286 (64%) were not previously mapped under physiological conditions (Figure 3B, right), and 70 were not found in proteomics screens under any previously investigated conditions (Table S4). Intriguingly, serine residues (55%) remained the most commonly ADP-ribosylated under physiological conditions (Figure 3B, left). Nonetheless, we also observed frequent modification of arginine (26%), histidine (12%), and cysteine (3.3%) residues. Physiological ADPr could be confidently localized (>90%) on cysteine, histidine, arginine, serine, threonine, and tyrosine residues (Figure 3C), while we did not observe consistent ADP-ribosylation of lysine, glutamate, and aspartate residues. We observed that under physiological conditions, >50% of serine ADPr resided in KS motifs (Figure S3A); an established phenomenon for serine ADPr (Larsen et al., 2018). Adherence to KS motifs was considerably lower in response to oxidative stress (Figure S3A), suggesting that under these conditions serine residues may be targeted more promiscuously. Physiological ADPr predominantly targeted nuclear proteins (71%; Figure S3B), although not to the same extent as H_2_O_2_-induced ADPr (85%; Figure S3C).

Next, we investigated the expression levels of proteins which have previously been observed to be ADP-ribosylated (Figure 3D). With the routine use of H_2_O_2_ induction of ADP-ribosylation, which greatly boosts the amount of ADPr in cells, we found that protein copy-numbers of ADPr target proteins span six orders of magnitude. We observed that physiologically ADP-ribosylated target proteins covered a comparable range of expression levels, although overall the depth of sequencing was somewhat lower. Nonetheless, compared to physiological ADPr target proteins identified in our previous study (Larsen et al., 2018), we achieved a 57% greater depth of sequencing (Figure 3D).

Overall, the extent of ADP-ribosylation was vastly different between unperturbed cells and H_2_O_2_-treated cells. Combining previous proteomics-based knowledge on ADPr (Bilan et al., 2017; Bonfiglio et al., 2017b; Hendriks et al., 2019; Larsen et al., 2018), with the additional insight gained in this study, we set out to investigate systemic differences between physiological and H_2_O_2_-perturbed ADPr biology. Specifically, we overlapped 9,029 ADPr sites mapped in response to H_2_O_2_ with 643 ADPr sites detected under physiological conditions (Figure 3E). The vast majority of ADPr sites (93%) were exclusive to H2O2 treatment, with 6% of ADPr sites detected in both cases, and 1% of ADPr sites exclusively detected under physiological conditions. In terms of amino acid distribution, the H_2_O_2_-exclusive ADPr predominantly modified serine residues (84%), in agreement with previous reports (Hendriks et al., 2019; Larsen et al., 2018; Palazzo et al., 2018). ADPr detected under all conditions still targeted a majority of serine residues (62%), but moreover modified significant fractions of arginine (13%), histidine (13%), cysteine (5%), and lysine (4%) residues. Intriguingly, physiologically exclusive ADPr primarily modified arginine residues (66%), with further modification of histidine (15%) and cysteine (10%) residues, and only a small fraction of modification on serine residues (5.7%). Still, our data revealed a propensity for ADPr to be targeted to different amino acid residues dependent on cellular condition.

Next, we performed functional annotation enrichment analysis to assess relative differences between physiological and non-physiological ADPr (Figure 3E and Table S5). H_2_O_2_-specific ADPr was more frequently observed to target proteins involved in rRNA-binding, DNA replication, and cell cycle, and less frequently modified glycosyltransferases, nucleosome proteins, and membrane proteins. Physiological-specific ADPr preferentially targeted proteins with sulfotransferase activity, glycoproteins, and membrane proteins, and averted targeting nuclear proteins involved in transcription, cell cycle, and the DNA damage response. Finally, proteins ADP-ribosylated under all conditions represented terms canonically associated with ADP-ribosylation, including predominant modification of the ADP enzymatic machinery, histone proteins, and DNA repair proteins, and this subset of ADPr target proteins less frequently included transcription factors, chromatin regulators, and proteins involved in cell cycle regulation. Collectively, our comparison between physiological and H_2_O_2_-induced ADPr demonstrates a considerable shift in the targeting of this PTM, not only at the substrate level, but also in terms of which amino acid residue types are modified.

### Detection of PARP8 and PARP14 auto-modification in cells

It is widely acknowledged that PARP1 modifies itself in response to DNA binding (Bolderson et al., 2019), and we profiled the main auto-modification of PARP1 to target three serine residues (Larsen et al., 2018). Our physiological profiling of ADP-ribosylation using AI-ETD corroborated that the same three serine residues are also modified under standard cellular growth conditions, with the majority (99.4%) of modification targeting Serine-499, followed by Serine-507 and Serine-519 (Figure 4A), with further physiological ADPr sites on PARP1 predominantly on serine residues.

**Figure 4.**
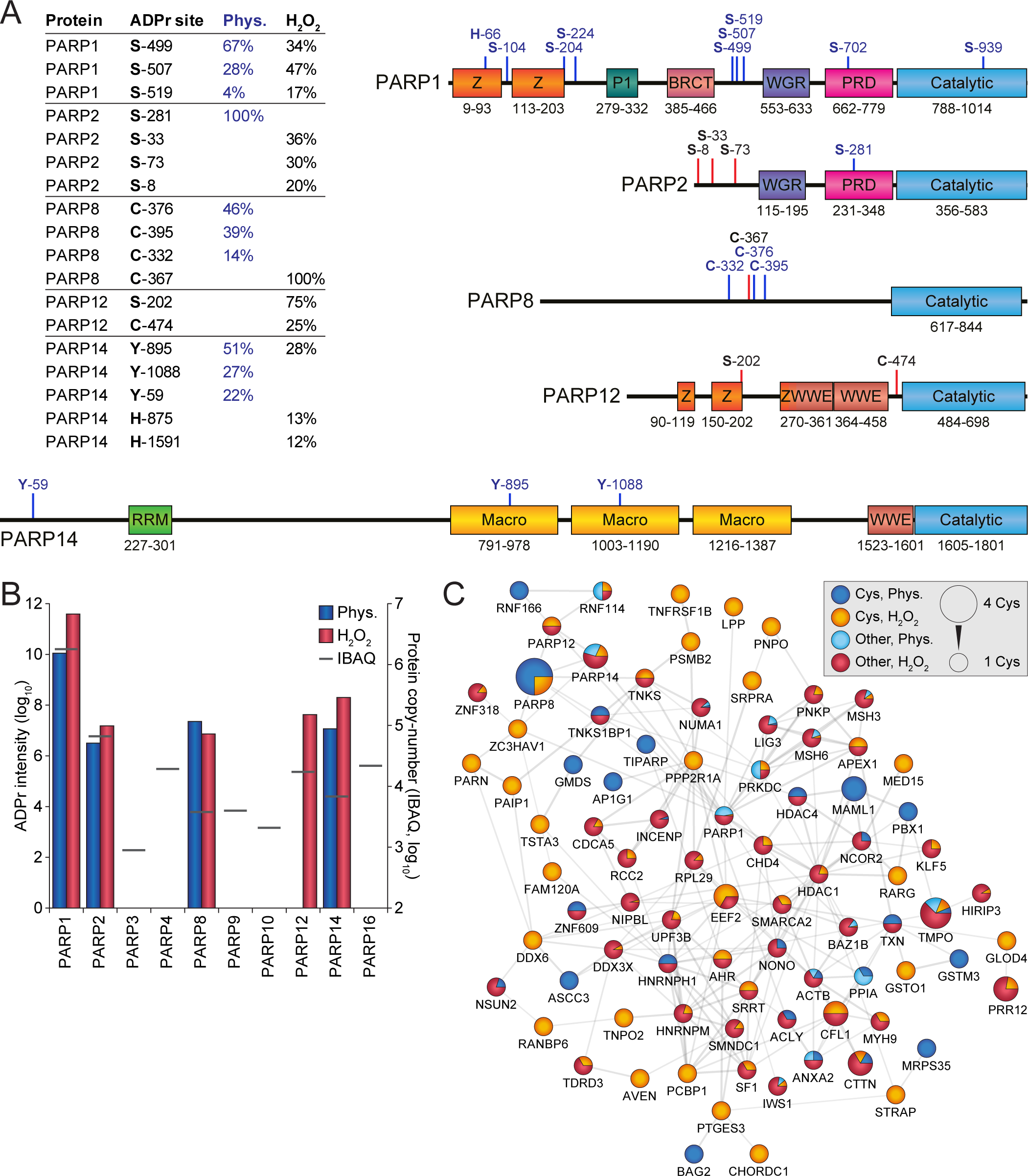
ADP-ribosylation of different PARP family members. (A) Overview of the most abundantly ADPr-modified residues within PARP family members, as a fraction of total protein ADPr modification, either under physiological conditions (“Phys.”) or in response to H_2_O_2_. All physiologically detected ADPr sites, or otherwise up to four of the most abundant total ADPr sites, are indicated on the graphical representation of the PARP family members. Blue lines; physiological ADPr sites, red lines; H_2_O_2_-induced ADPr sites. (B) Overview of the total amount of ADPr detected on each PARP family member, in relation to their known expression level (“IBAQ”). Both values are logarithmic, and graph scaling was normalized to PARP1 and PARP2 values. (C) STRING network visualizing functional interactions between proteins identified to be ADPr-modified on cysteine residues in this study or in Larsen et al., 2018 (Larsen et al., 2018). Default STRING clustering confidence was used (*P*>0.4), and disconnected proteins were omitted from the network. Proteins were significantly interconnected, with a protein-protein interaction enrichment *q*-value of 1.1*10^−11^. Nodes corresponding to proteins were annotated with visual properties, as highlighted in the figure legend. “Cys”; modification on cysteine residues, “Other”; modification on non-cysteine residues, “Phys.”; physiological conditions. Distribution of color is relative to the number of sites within each category, sites detected both physiologically and in response to H_2_O_2_ were colored as physiological.

With the improved analytical sensitivity gained by AI-ETD, we were curious whether we could detect ADPr modification of other PARP enzymes, under physiological conditions or otherwise in response to H_2_O_2_ treatment. Based on deep proteomic profiling of HeLa cells, ten PARP family members are expressed at different levels in unperturbed HeLa cells (Bekker-Jensen et al., 2017). Using AI-ETD, we could detect ADPr modification of PARP1, PARP2, PARP8, PARP12, and PARP14 (Figure 4B and Tables S3 and S4). Strikingly, even though protein copy-numbers of PARP8 and PARP14 are 250 to 500 times lower than PARP1, we could still detect their ADP-ribosylation under physiological conditions.

On PARP2, we only observed modification of serine residues, similar to PARP1 (Figure 4A), corroborating the known inter-connected functions of the two enzymes (Menissier de Murcia et al., 2003; Schreiber et al., 2002). ADP-ribosylation on PARP12 was only found in response to H_2_O_2_ treatment, and at relatively low abundance. Intriguingly, PARP8 was ADP-ribosylated on three cysteine residues under physiological conditions, Cysteine-376, Cysteine-395, and Cysteine-332, whereas a fourth cysteine residue, Cysteine C-367, was modified in response to H_2_O_2_. This exclusive modification on cysteine residues could hint at PARP8 auto-modification, which may imply that PARP8 is capable of ADP-ribosylation of cysteine residues. Analogously, we observed physiological modification of three tyrosine residues in PARP14, Tyosine-895, Tyrosine-1088, and Tyrosine-59, which could likewise imply auto-modification, and a potential function of PARP14 as the catalyst for tyrosine ADP-ribosylation.

ADP-ribosylation of serine, arginine, tyrosine, and histidine residues, has previously been reported (Hendriks et al., 2019). We were intrigued by the ADPr modification of cysteine residues, with PARP8 modified on four cysteine residues, and knowledge regarding cysteine ADPr being relatively sparse. Altogether, we could map 123 cysteine ADPr sites to 112 proteins (Tables S3 and S4), with only one cysteine ADPr site detected for the majority of proteins. We generated an interconnected STRING network based on all proteins with cysteine ADPr (Figure 4C), with modification occurring under physiological conditions or in response to oxidative stress. 85 out of 112 proteins (76%) were connected to the network, with significant enrichment for protein-protein interactions. Beyond PARP8, cysteine ADPr modification occurred on a considerable number of interconnected catalytic ADPr enzymes, including PARP1, PARP5A (TNKS), PARP7 (TIPARP), PARP12, PARP13 (ZC3HAV1), and PARP14. When only considering the 60 proteins predominantly ADP-ribosylated on cysteines, there was a significant presence of proteins involved in RNA metabolism, including DDX6, HNRNPH1, NSUN2, PAIP1, PARN, PCBP1, PPP2R1A, PSMB2, TNKS1BP1, and TRNT1. Taken together, our sensitive MS-based proteomics screening of ADPr provides further insight into modification of non-serine residue types, and may hint at enzymatic specificity of certain PARP enzymes towards specific amino acids.

## DISCUSSION

Here, we investigated the potential of AI-ETD for sensitive identification and confident localization of ADPr sites. To evaluate the ability of AI-ETD for profiling the ADP-ribosylome, we performed a comparison of pure ETD, EThcD, and AI-ETD at five different laser power settings.

ADP-ribosylated peptides are very labile when subjecting them to higher-energy collisional dissociation (HCD), with the ADP-ribose itself drawing much of the energy and being one of the first fragmentation events, ultimately prohibiting localization of the ADP-ribose to the correct amino acid, as no fragment ions remain with the ADP-ribose attached (Bonfiglio et al., 2017a; Larsen et al., 2018). Contrarily, ADP-ribosylated peptides are very stable when subjected to ETD, and here we observed ADPr peptides to be much more prone to non-dissociative electron transfer (ETnoD), compared to unmodified peptides within the same samples. Supplemental activation of the ETD reaction with HCD, i.e. EThcD, still largely results in an ETD-type fragmentation, while somewhat mitigating ETnoD and generating more comprehensive peptide bond coverage (Bilan et al., 2017; Hendriks et al., 2019). However, EThcD fails to completely overcome ETnoD, and results in decreased spectral quality at higher energies. Intriguingly, we observed that ADPr peptides are highly susceptible to AI-ETD, much more so than small unmodified peptides within the same sample. This susceptibility to AI-ETD resulted in a near-complete abolishing of ETnoD at laser power settings of 25% and higher, although these greater energies also resulted in over-fragmentation of the peptides and a sharp drop in spectral quality. At a modest level of energy, AI-ETD resulted in higher quality spectra, improved sensitivity, and more confident localization of ADP-ribosylation to acceptor amino acids, overall doubling the number of identifications compared to pure ETD.

Recent advances in mass spectrometry-based proteomics have largely focused on high-throughput analyses, essentially by minimizing time or input material required to achieve the same depth of sequencing (Bache et al., 2018; Batth et al., 2019). Compared to relatively abundant post-translational modifications such as phosphorylation and glycosylation, for which MS methodology is well-established (Hogrebe et al., 2018; Kelstrup et al., 2018; Riley et al., 2019), the study of ADP-ribosylation with MS-based proteomics remains daunting, requiring large amounts of input material and significant investments of mass spectrometric machine time (Hendriks et al., 2019; Larsen et al., 2018; Leutert et al., 2018; Li et al., 2019; Martello et al., 2016; Zhang et al., 2013). Using AI-ETD, we were able to identify >500 ADPr sites from single shot analyses of purified ADPr material originating from only ∼5 million cells. Moreover, from fractionated samples originating from ∼100 million cells, we were able to identify >5,000 ADPr sites, reaching a great depth of sequencing while using considerably less starting material than other studies (Hendriks et al., 2019; Larsen et al., 2018; Leutert et al., 2018; Li et al., 2019; Martello et al., 2016; Zhang et al., 2013).

Compared to ADPr induced by oxidative stress, the identification of ADPr in unperturbed cells is exceptionally challenging, with the modification occurring at very low abundance and dynamically regulated (Daniels et al., 2015). We previously reported a few hundred ADPr sites under physiological conditions, without the need for genetic perturbation of eraser enzymes, and without exposing cells to high levels of genotoxic stress (Larsen et al., 2018). However, this previous identification of physiological ADPr sites required an extensive investment of mass spectrometric machine time and a larger amount of starting material, with a considerable portion of identified ADPr sites relying on pre-fractionation of samples prior to mass spectrometric analysis. With the increased analytical sensitivity of AI-ETD, we were now able to map 450 physiological ADPr sites from single shot analyses, while using limited starting material and a total of 200 minutes of analytical gradient time for four replicates. Taken together, we demonstrate that AI-ETD can assist the study of physiological ADPr under unperturbed conditions, ultimately reducing the amount of starting material required for sequencing the native ADP-ribosylome, and facilitating studying ADP-ribosylation in a biomedical context.

Auto-modification of PARPs is a known phenomenon (Vyas et al., 2014), with especially auto-modification of PARP1 being thoroughly studied (Chapman et al., 2013; Muthurajan et al., 2014). Here, we were able to identify ADPr occurring on PARP1, 2, 8, 12, and 14. For PARP1 and PARP2, serine was the main acceptor residue. This is consistent with PARP1 and PARP2 catalyzing ADPr of serine residues upon oxidative stress, in combination with HPF1 (Bonfiglio et al., 2017b; Larsen et al., 2018; Palazzo et al., 2018). While we observed a relatively high number of serine residues being ADP-ribosylated under physiological conditions, many of those were also detectable in response to oxidative stress. Contrarily, ADPr sites exclusively detected under physiological conditions were only rarely modifying serine residues. Taken together, this would suggest that serine ADP-ribosylation occurs predominantly in response to DNA damage, and that serine ADP-ribosylation observed under physiological conditions could be a result of baseline DNA damage, potentially as a consequence of culturing cells under atmospheric oxygen conditions.

Intriguingly, we only observed PARP8 to be ADP-ribosylated on cysteine residues. This exclusive modification of cysteine residues supports previous *in vitro* observations of the auto-modification of PARP8 (Vyas et al., 2014), and we confirmed all four hitherto reported cysteine residues, of which three were detectable under physiological conditions. In case of PARP14, physiological ADP-ribosylation only targeted tyrosine residues, similarly setting it apart from PARP1 and PARP2 which were primarily modified on serine residues. Auto-modification of PARP14 has been described previously (Qin et al., 2019), and the interactome of PARP14 suggests a role in regulating RNA stability (Carter-O’Connell et al., 2018). We detected modification of PARP13 in cells on one cysteine residue, and PARP14 has been demonstrated to MARylate PARP13 *in vitro* (Carter-O’Connell et al., 2018), although there acidic amino acids were targeted for modification. Overall, the auto-modification of PARP8 and PARP14 could suggest that these enzymes may have the ability to specifically ADP-ribosylate cysteine and tyrosine residues, respectively. However, further investigation is required to elucidate whether, and how, such specificity would be mediated.

Taken together, the combination of the AI-ETD technology with our Af1521 strategy for unbiased purification of ADP-ribosylated peptides, has culminated in a highly sensitive MS-based proteomics approach that can confidently identify large numbers of ADP-ribosylation events from limited starting material. Moreover, this increased sensitivity makes it feasible to profile ADP-ribosylation in an entirely physiological context, taking a final step towards biomedical and clinical applicability.

## AUTHOR CONTRIBUTIONS

S.C.L. and I.A.H. prepared all samples. M.W., J.M.L, E.S., and J.J.C. established AI-ETD technology and infrastructure. S.C.L., I.A.H., and J.M.L. measured all samples on the mass spectrometer. I.A.H. and S.C.L. processed all MS raw data. I.A.H., S.C.L., and M.R. performed bioinformatics and statistical analyses. M.R. and B.F. wrote the Python script for determination of fragmentation efficiency. M.L.N. and J.J.C. supervised the project. I.A.H., S.C.L., and M.L.N. wrote the manuscript with input from all authors.

## ACKNOWLEDGMENTS

The work carried out in this study was in part supported by the Novo Nordisk Foundation Center for Protein Research, the Novo Nordisk Foundation (grant agreement numbers NNF14CC0001 and NNF13OC0006477), The Danish Council of Independent Research (grant agreement numbers 4002-00051, 4183-00322A and 8020-00220B), and The Danish Cancer Society (grant agreement R146-A9159-16-S2). The proteomics technology applied were part of a project that has received funding from the European Union’s Horizon 2020 research and innovation program under grant agreement EPIC-XS-823839. We would like to acknowledge the lab of Michael O. Hottiger for the expression and purification of recombinant human PARG (University of Zurich), and we thank members of the NNF-CPR Mass Spectrometry Platform for instrument support and technical assistance.

## METHODS

### Cell culture

HeLa cells (CCL-2) were acquired via the American Type Culture Collection, and cultured at 37 °C and 5% CO2 in Dulbecco’s Modified Eagle’s Medium (Invitrogen) supplemented with 10% fetal bovine serum and a penicillin/streptomycin mixture (100 U/mL; Gibco). Cells were routinely tested for mycoplasma. Cells were not routinely authenticated. For initial optimization and benchmarking experiments, ADP-ribosylation was induced in HeLa by treatment of the cells with 1 mM H_2_O_2_ (Sigma Aldrich) for 10 min in PBS at 37°C. Five cell culture replicates were prepared, each corresponding to approximately 100 million HeLa cells, of which one was used for optimization, and four for benchmarking. For physiological experiments, cells were left untreated, and four cell culture replicates were prepared, each corresponding to approximately 100 million HeLa cells.

### Cell lysis and protein digestion

The full procedure for enrichment of ADPr from cells was done as described previously (Hendriks et al., 2019; Larsen et al., 2018). Briefly, cells were washed twice with ice-cold PBS, and gently scraped at 4 °C in a minimal volume of PBS. Cells were pelleted by centrifugation at 500*g*, and lysed in 10 pellet volumes of Lysis Buffer (6 M guanidine-HCl, 50 mM TRIS, pH 8.5). Complete lysis was achieved by alternating vigorous shaking with vigorous vortexing, for 30 seconds, prior to snap freezing of the lysates using liquid nitrogen. Frozen lysates were stored at −80 °C until further processing. Lysates were thawed and sonicated at 30 W, for 1 second per 1 mL of lysate, spread across 3 separate pulses. Tris(2-carboxyethyl)phosphine and chloroacetamide were added to a final concentration of 5 mM, and the lysate was incubated for 1 hour at 30 °C. Proteins were digested using Lysyl Endopeptidase (Lys-C, 1:100 w/w; Wako Chemicals) for 3 hours, and diluted with three volumes of 50 mM ammonium bicarbonate. Half of the samples was further digested overnight using modified sequencing grade Trypsin (1:200 w/w; Sigma Aldrich), and the other half was re-digested overnight using Lys-C. Digested samples were acidified by addition of trifluoroacetic acid (TFA) to a final concentration of 0.5% (v/v), cleared by centrifugation, and purified using reversed-phase C18 cartridges (SepPak Classic, 360 mg sorbent, Waters) according to the manufacturer’s instructions. Elution of peptides was performed with 30% or 40% ACN in 0.1% TFA, for peptides digested with either trypsin or Lys-C, respectively. Eluted peptides were frozen overnight at −80 °C, and lyophilized.

### Purification of ADP-ribosylated peptides

GST-tagged Af1521 macrodomain was produced in-house and coupled to glutathione Sepharose 4B beads (Sigma-Aldrich), essentially as described previously (Hendriks et al., 2019; Larsen et al., 2018). Lyophilized peptides were dissolved in IP buffer (50 mM TRIS pH 8.0, 1 mM MgCl_2_, 250 μM DTT, and 50 mM NaCl), after which long ADP-ribose polymers were reduced to monomers using recombinant PARG at a concentration of 1:10,000 (w/w), overnight and at room temperature. Subsequently, 100 μL of sepharose beads with GST-tagged Af1521 were added to the samples, and mixed at 4 °C for 3 h. Beads were sequentially washed four times with ice-cold IP Buffer, two times with ice-cold PBS, and two times with ice-cold MQ water. On the first wash, beads were transferred to 1.5 mL LoBind tubes (Eppendorf), and LoBind tubes were exclusively used from this point on to minimize loss of peptide. Additional tube changes were performed every second washing step to minimize carryover of background. ADP-ribosylated peptides were removed from the beads using two elution steps with 200 μL ice-cold 0.15% TFA, and the pooled elutions were cleared through 0.45 μm spin filters (Ultrafree-MC, Millipore) and subsequently through pre-washed 100 kDa cut-off filters (Vivacon 500, Sartorius). The filtered ADP-ribosylated peptides were purified using C18 StageTips at high pH (Hendriks et al., 2019), and eluted either as single shot samples (optimization and physiological experiments), or eluted as seven fractions (benchmarking experiments). Briefly, samples were basified by adding ammonium solution to a final concentration of 20 mM, and loaded onto StageTips carrying four layers of C18 disc material (punch-outs from 47mm C18 3M™ extraction discs, Empore). Elution was performed with 25% of acetonitrile (ACN) in 20 mM ammonium hydroxide for single shot samples, or sequentially performed with 2% (F1), 4% (F2), 7% (F3), 10% (F4), 15% (F5), and 25% (F6) of ACN in 20 mM ammonium hydroxide for fractionated samples. The seventh fraction (F0) was prepared by performing StageTip purification at low pH on the flowthrough fraction from sample loading at high pH. All samples were completely dried using a SpeedVac at 60 °C, and dissolved in a small volume of 0.1% formic acid. Final samples were frozen at −20 °C until measurement.

### Mass spectrometric analysis

All MS samples were analyzed using a Fusion Lumos Orbitrap mass spectrometer (Thermo Fisher Scientific, San Jose, CA, USA) modified for AI-ETD, coupled to a Dionex UltiMate 3000 nano-UHPLC system (Thermo Fisher, Sunnyvale, CA, USA). For AI-ETD implementation, a Firestar T-100 Synrad 60 Watt (W) CO_2_ continuous wave laser (Mukilteo, WA, USA) was mounted at the back of the instrument. The IR photon beam was guided into the traps using focusing lenses and a multi-mode hollow-core fiber (Opto-Knowledge Systems, Torrance, CA, USA) prior to a zinc selenide window that was placed in the vacuum manifold. Chromatographic separation of peptides was performed on a 30-cm long analytical column, with an internal diameter of 1.7 μm, packed in-house as described previously (Shishkova et al., 2018) using 130 Å pore size Bridged Ethylene Hybrid C18 particles (Waters, Milford, USA). The analytical column was heated to 50 °C using an in-house made column oven. Peptides were eluted from the column using a gradient of Buffer A (0.1% formic acid) and Buffer B (80% ACN in 0.1% formic acid). The primary gradient ranged from 3% buffer B to 24% buffer B over 50 minutes, preceded by sample loading time (5 minutes for evaluation experiments, 20 minutes for benchmarking and physiological experiments), and followed by a 20 minute washing block. Electrospray ionization (ESI) was achieved using a Nanospray Flex Ion Source (Thermo). Spray voltage was set to 2.25 kV, capillary temperature to 275°C, and RF level to 30%. Full scans were performed at a resolution of 60,000, with a scan range of 300 to 1,750 m/z, a maximum injection time of 60 ms, and an automatic gain control (AGC) target of 600,000 charges. Precursors were isolated with a width of 1.3 m/z, with an AGC target of 200,000 charges, and precursor fragmentation was accomplished using either electron transfer dissociation (ETD), electron transfer disassociation with supplemental higher-collisional disassociation (EThcD) at an NCE of 20, or activated-ion electron transfer dissociation (AI-ETD); all using calibrated charge-dependent ETD parameters (Rose et al., 2015). The laser powers were the percentage of power (W) from the Firestar T-100 Synrad 60-W CO_2_, continuous wave laser. For AI-ETD evaluation, laser power settings of 10% (6 W), 15% (9 W), 20% (12 W), 25% (15 W), and 30% (18 W) were used, and for benchmarking and physiological experiments a laser power setting of 20% was used. Precursors with charge state 2-5 were isolated for MS/MS analysis, and prioritized from charge 3 (highest), to 4, to 5, to 2 (lowest), using the decision tree algorithm. For benchmarking and physiological experiments, only precursors with charge state 3-5 were isolated. Precursors were excluded from repeated sequencing by setting a dynamic exclusion time of 75 seconds for evaluation experiments, and 90 seconds for benchmarking and physiological experiments, with an exclusion mass tolerance of 10 ppm. MS/MS spectra were measured in the Orbitrap, with 5 data-dependent MS/MS scans per full MS scan, a maximum precursor injection time of 120 ms, and a scan resolution of 60,000. First mass for MS/MS scans was set to 100 for ETD and EThcD measurements, and variable for AI-ETD measurements, with the first mass defining the laser power applied. MS/MS first masses for AI-ETD were 141, 131, 142, 132, and 143, corresponding to application of laser powers 10%, 15%, 20%, 25%, and 30%, respectively.

### Data analysis

Analysis of the mass spectrometry raw data was performed using MaxQuant software (version 1.5.3.30). MaxQuant default settings were used, with exceptions outlined below. Two separate computational searches were performed, one for evaluation and physiological data, and the other for benchmarking data. For generation of the theoretical spectral library, a HUMAN.fasta database was extracted from UniProt on the 24^th^ of May, 2019. N-terminal acetylation, methionine oxidation, cysteine carbamidomethylation, and ADP-ribosylation on all amino acid residues known to potentially be modified (C, D, E, H, K, R, S, T, and Y), were included as variable modifications. For analysis of trypsin samples, up to 6 missed cleavages were allowed. For analysis of Lys-C samples, up to 3 missed cleavages were allowed. A maximum allowance of 4 variable modifications per peptide was used. Second peptide search was enabled (default), and matching between runs was enabled with a match time window of 42 seconds and an alignment time window of 20 minutes. Mass tolerance for precursors was set to 20 ppm in the first MS/MS search and 4.5 ppm in the main MS/MS search after mass recalibration. For fragment ion masses, a tolerance of 20 ppm was used. Modified peptides were filtered to have an Andromeda score of >40 (default), and a delta score of >20. Data was automatically filtered by posterior error probability to achieve a false discovery rate of <1% (default), at the peptide-spectrum match, the protein assignment, and the site-specific levels.

### Data filtering

Beyond automatic filtering and FDR control as applied by MaxQuant, the data was manually filtered in order to ensure proper identification and localization of ADP-ribose. PSMs modified by more than one ADP-ribose were omitted. PSMs corresponding to unique peptides were only used for ADP-ribosylation site assignment if localization probability was >0.90, with localization of >0.75 accepted only for purposes of intensity assignment of further evidences. Erroneous MaxQuant intensity assignments were manually corrected in the sites table, and based on localized PSMs only (>0.90 best-case, >0.75 for further evidences). For the ADP-ribosylation target proteins table, the proteinGroups.txt file generated by MaxQuant was filtered to only contain those proteins with at least one ADP-ribosylation site detected and localized post-filtering as outlined above, with cumulative ADP-ribosylation site intensities based only on localized evidences.

### Determination of fragmentation efficiency

An in-house Python script was used to determine the intensities of non-reduced precursors (no ETD), and the corresponding non-dissociated charge-reduced precursor ions (ETnoD) for each MS/MS spectrum. For that, the centroided, de-isotoped, and charge-deconvoluted peak lists from the MaxQuant Andromeda Peak List (APL) files were used. Second peptide search APLs were not considered. “No ETD” precursor peaks were expected at:

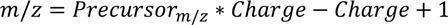

“ETnoD” peaks were expected at:

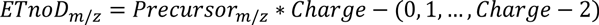

This compensates for MaxQuant charge deconvolution, which goes by the assumption that the charge state is purely dependent on added protons. The expected peaks were matched to peaks from the MaxQuant APL files, with a mass tolerance of 0.05 Da, and their intensities were stored. Intensity information was mapped to the msms.txt output file from MaxQuant using the scan number. Total fragmentation of peptides was calculated by division of fragment product ion intensities with the sum of all peak intensities.

### Comparison to other studies

Data from other studies was retrieved from several publications and online databases. For ADP-ribosylation sites; Larsen et al. (Larsen et al., 2018), Hendriks et al. (Hendriks et al., 2019), Bonfiglio et al. (Bonfiglio et al., 2017b), Bilan et al. (Bilan et al., 2017), Leslie Pedrioli et al. (Leslie Pedrioli et al., 2018). For ADP-ribosylation proteins; Zhang et al. (Zhang et al., 2013), Jungmichel et al. (Jungmichel et al., 2013), Martello et al. (Martello et al., 2016), Bilan et al. (Bilan et al., 2017), Larsen et al. (Larsen et al., 2018), Hendriks et al. (Hendriks et al., 2019). For total proteome; Bekker-Jensen et al. (Bekker-Jensen et al., 2017). For comparison of proteins between studies, all protein identifiers were mapped to the human proteome as downloaded from Uniprot on the 24th of May, 2019. In case data sources did not include Uniprot IDs, ID mapping on Uniprot was used to convert other IDs to Uniprot IDs, and otherwise gene names were used. Non-existent, non-human, and redundant entries, were discarded from the analysis. For comparison of sites between studies, all parent proteins were mapped to Uniprot IDs as described above, and afterwards the reported positions of modification were used to extract 51 amino acid sequence windows (modified residue +/− 25 amino acids), with the sequence windows ultimately used to directly compare identified sites. Sites mapping to non-existent, non-human, or redundant proteins, were discarded. Sites not correctly aligning to the reported amino acid residues were discarded.

### Statistical analysis

Details regarding the statistical analysis can be found in the respective figure legends. Statistical handling of the data was primarily performed using the freely available Perseus software (Tyanova et al., 2016), and includes term enrichment analysis through FDR-controlled Fisher Exact testing, and density estimation for highly populated scatter plots. Protein Gene Ontology annotations and UniProt keywords used for term enrichment analysis were concomitantly downloaded from UniProt with the HUMAN.fasta file used for searching the RAW data. Boxplots and violin plots were generated using the BoxPlotR web tool (Spitzer et al., 2014). The online STRING database (version 11) was used for generation of protein interaction networks (Szklarczyk et al., 2017), and Cytoscape (version 3.7.1) was used for manual annotation and visualization of the STRING networks (Shannon et al., 2003).

---

**Figure S1.**
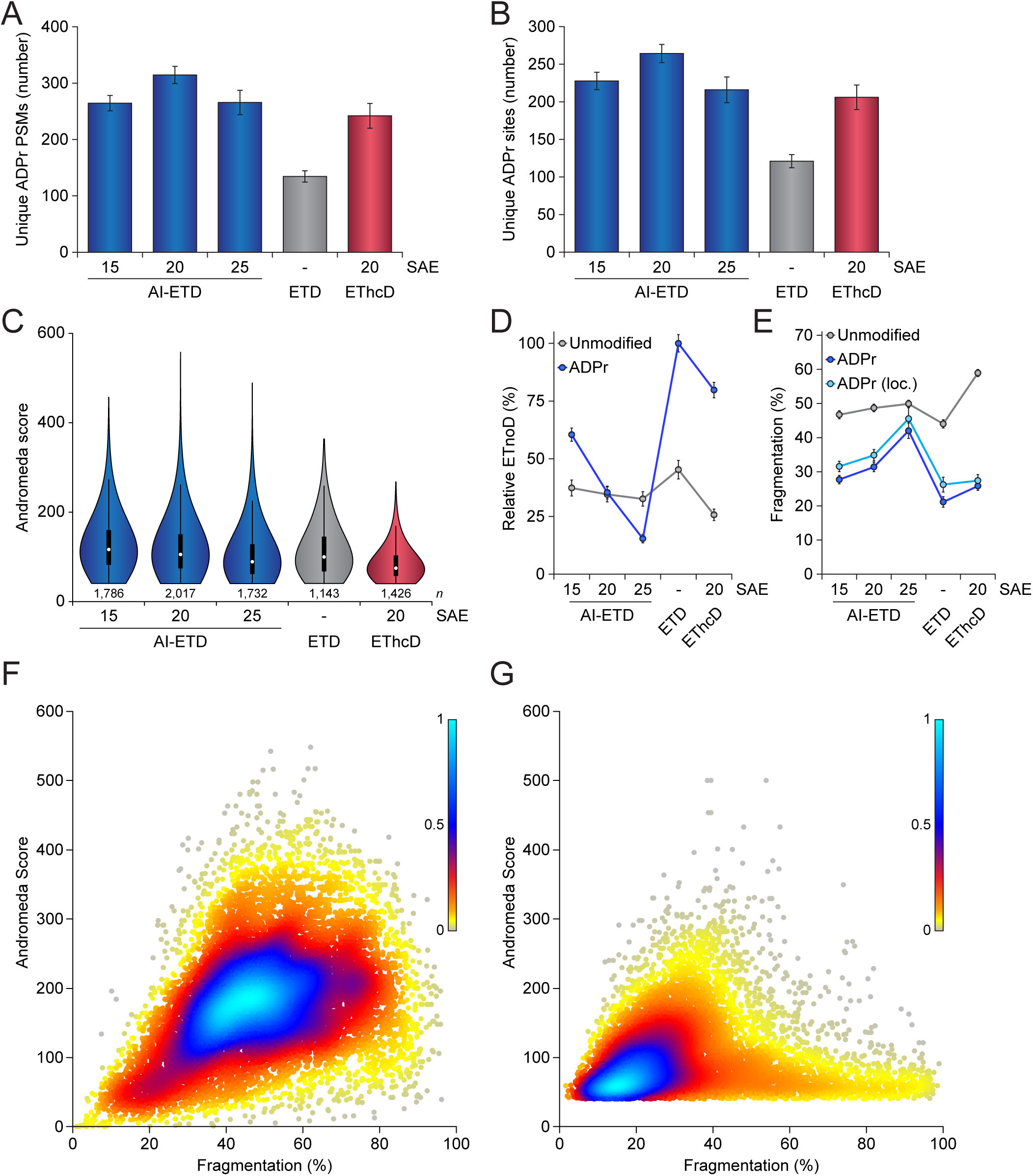
Comparison of AI-ETD vs. ETD and EThcD for mapping ADP-ribosylation in Lys-C digested peptides. (A) Overview of the number of ADPr peptide-spectrum-matches (PSMs) identified and localized (>90% probability) for each dissociation method and Supplemental Activation Energy (SAE). Error bars represent SD, *n*=4 technical replicates. (B) As **A**, but displaying the number of ADPr sites identified. (C) As **A**, but displaying the spectral quality (in Andromeda Score) of all identified ADPr-modified peptides. Distribution of data points is visualized, line limits; 1.5× interquartile range (IQR), box limits; 3^rd^ and 1^st^ quartiles, white dot; mean. Number of data points (*n*) is visualized below the distributions. (D) Visualization of the average relative degree of non-dissociative electron transfer (ETnoD). Derived from all peptide-identified MS/MS spectra, and separately visualized for unmodified and ADP-ribosylated peptides. Error bars represent 5× SEM. (E) Visualization of the average degree of precursor fragmentation, calculated by dividing observed fragment ion peak intensity by the sum of non-ETD, ETnoD, and all fragment ion peak intensities. Derived from all peptide-identified MS/MS spectra, and separately visualized for unmodified, ADP-ribosylated, and localized ADP-ribosylated peptides. Error bars represent 5× SEM. (F) Spectral quality (in Andromeda Score) plotted against the average degree of precursor fragmentation, for unmodified peptides detected within ADPr samples. Coloring represents the relative density of dots in the plot, with higher values corresponding to higher density. (G) As **F**, but for non-localized ADP-ribosylated peptides.

**Figure S2.**
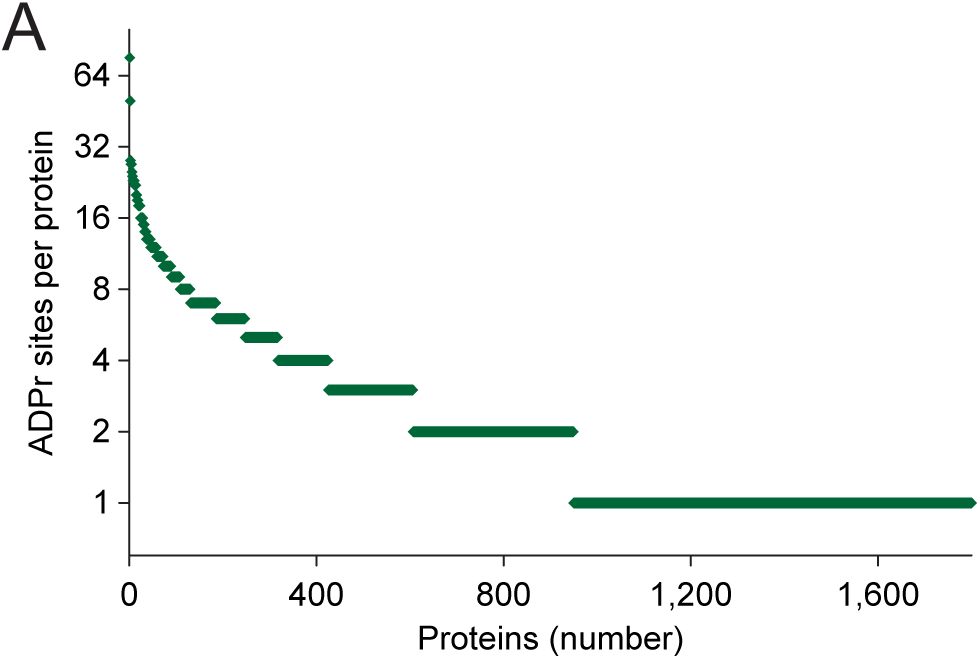
Distribution of ADPr sites across proteins. (A) Visualization of the number of unique ADPr sites detected per protein.

**Figure S3.**
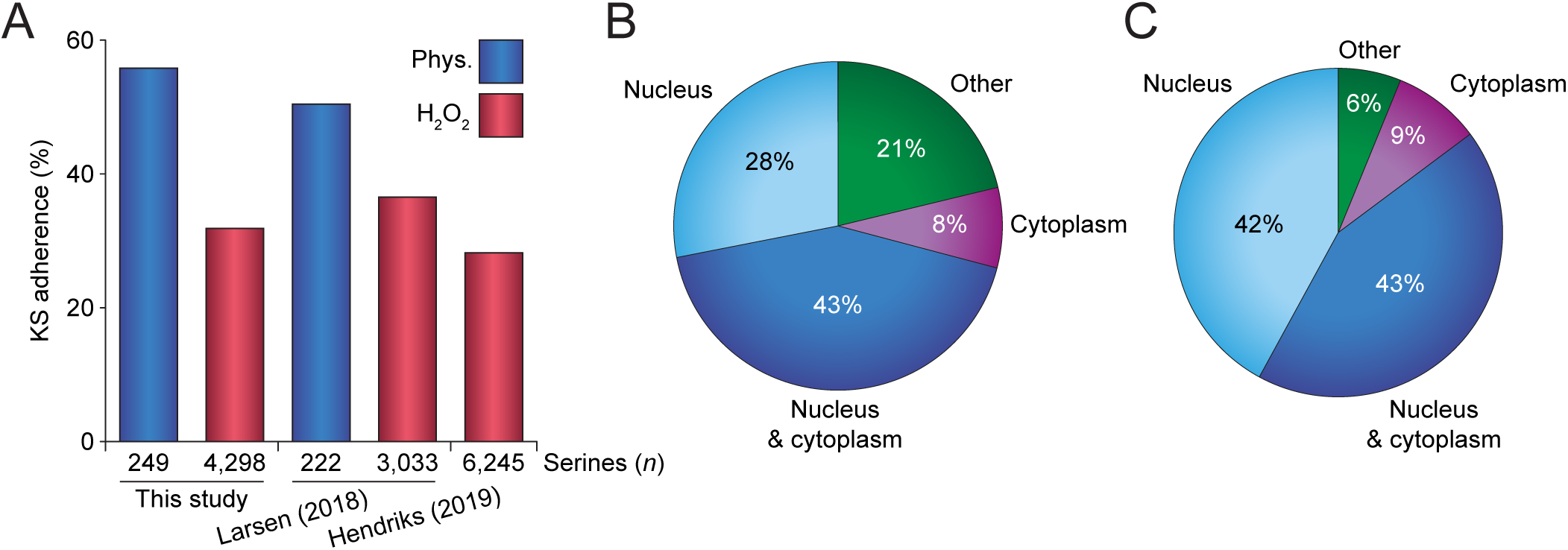
KS motif adherence and subcellular localization. (A) Overview of KS motif adherence of all serine ADPr sites identified under physiological conditions or in response to H_2_O_2_ treatment, in this study or in two others (Hendriks et al., 2019; Larsen et al., 2018). KS motif modification is defined as an ADPr-modified serine with an N-terminal lysine residue. (B) Subcellular localization of ADPr target proteins identified under physiological conditions, with localization derived from Gene Ontology Cellular Compartments. (C) As **B**, but for ADPr target proteins identified in response to H_2_O_2_ treatment.

